# Asymmetric Structural Transfer Between Natural Language and Biological Foundation Models

**DOI:** 10.64898/2026.07.09.737502

**Authors:** Liang Wang

**Affiliations:** School of Artificial Intelligence and Automation, Huazhong University of Science and Technology, Wuhan 430070, P.R. China

## Abstract

Cross-domain transfer is a defining property of foundation models, yet whether such transfer is *symmetric* across domains remains unknown. Prior work has reported a striking transfer from natural language to biological sequences: language models fine-tuned only on English structural tasks acquire zero-shot protein-homology discrimination. Here we ask the converse and general question—*is structural transfer between language and biology directional?* —and answer it systematically. We first reproduce forward transfer (language →biology) under controlled conditions, then evaluate the reverse direction (biology →language) across fine-tuning, iso-token continued pretraining, model scaling, multiple biological foundation-model families (ESM-2, ProtBERT), and adversarial synthetic structure tasks. Reverse transfer is consistently weak: it does not exceed matched-token controls, does not scale, and does not generalize. In an architecture-matched 2 *×* 2 analysis on models with *known* training data—which eliminates the pretraining-contamination confound that clouds large-model studies—a language model retains far more competence when moved to biology (off-domain drop 0.08) than a biological model retains when moved to language (drop 0.36). Scaling widens rather than closes this gap: language →biology transfer strengthens with size while biology →language transfer decays toward chance, a pattern shared by two independent protein-model families. Our findings establish that shared structural regularities between natural language and biological sequences do *not* imply symmetric representational transfer, revealing an intrinsic directionality in cross-domain foundation-model learning.

## 1 Introduction

The emergence of foundation models [1, 2, 3] has made *cross-domain transfer* —the reuse of representations learned in one domain to solve tasks in another [4, 5, 6]—one of their most consequential and least understood properties. It underlies transfer across languages [7, 8], across modalities, and even into specialized scientific domains [9]. A particularly surprising instance has been reported between human language and the “language of life”: models optimized purely on natural-language structure acquire, without biological pretraining, a latent ability to discriminate homologous biological sequences [10, 11, 12]. This observation invites a bold interpretation—that abstract linguistic structure constitutes a universal cognitive prior for decoding biology, echoing the emergent, hard-to-predict capabilities documented in large models [13].

Such a claim, however, raises an immediate and rarely examined question. If natural language and biological sequences truly share a common structural substrate, then transfer between them should be a property of that shared substrate—and one would expect it to operate in *both* directions. Yet essentially all evidence to date concerns a single direction (language → biology), and much of it relies on large proprietary models whose training corpora may already contain biological data— large models are known to memorize training content [14]—confounding genuine structural transfer with memorization [15, 16]. Protein and genomic foundation models [17, 18, 19, 20, 21, 22] and structure predictors [23] are typically trained on domain data from scratch, leaving open whether their competence could instead arise from generic sequence structure. Whether the reverse transfer exists, and whether cross-domain structural transfer is symmetric at all, is unknown.

Here we treat directionality itself as the object of study. We ask: *is structural transfer between natural language and biological sequences symmetric, and if not, what governs its direction?* We answer through a systematic bidirectional analysis. (i) We reproduce forward transfer under controlled, multi-seed conditions on models with known training data (Part I). (ii) We evaluate reverse transfer (biology → language) exhaustively—classification fine-tuning across 100 seeds, iso-token continued-pretraining ablations with shuffled and random controls, model scaling, two biological foundation-model families, and an adversarial nested-structure task (Part II). (iii) We quantify the asymmetry directly on architecture-matched models whose training data is fully known, eliminating the contamination confound (Part III). (iv) We probe its mechanistic origin (Part IV).

Our central finding is that transfer is *bidirectional but strongly and universally asymmetric* (Fig. 1). Language→biology transfer is robust and strengthens with scale; biology→language transfer is weak, does not exceed matched-token controls, and *decays* toward chance as biological models grow—a pattern reproduced across ESM-2 and ProtBERT. Mechanistic analysis localizes the asymmetry not to static representational alignment or an induced attention circuit, but to learnability: biological discrimination emerges as a by-product of learning language structure, but not the converse. We conclude that shared structural regularity does not imply symmetric transfer, and that natural language functions as the more general structural prior. This directionality, rather than any single transfer result, is the contribution: it reframes “can models cross domains?” as “why is crossing easier in one direction?”—a question about the nature of structure in foundation models.

**Figure 1:**
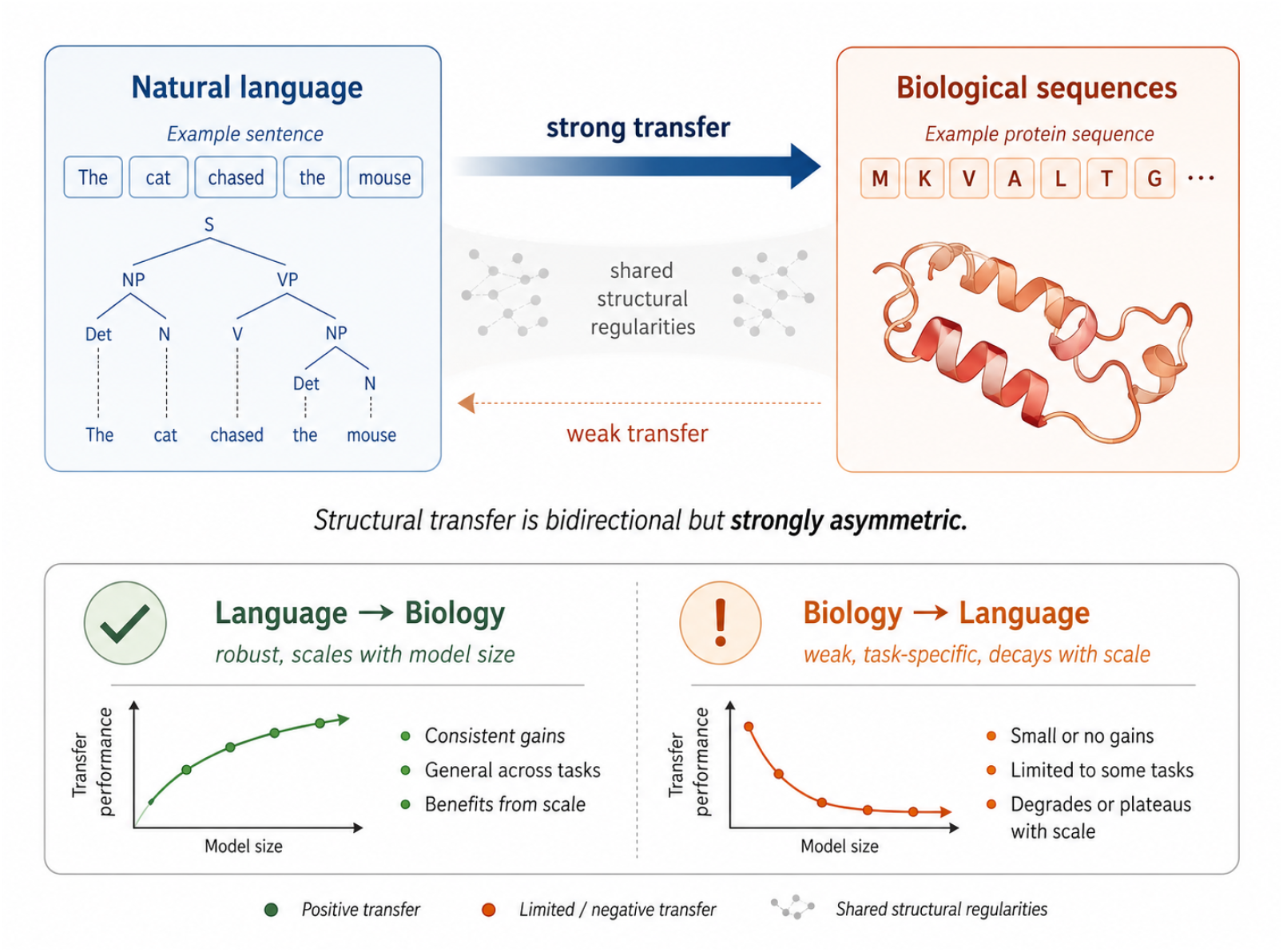
Structural transfer between natural language and biological sequences is directional. Forward transfer (language →biology) is robust and strengthens with model scale (off-domain competence drop 0.08); reverse transfer (biology →language) is weak, does not exceed matched-token controls, and decays with scale (drop 0.36). Transfer is thus bidirectional but strongly asymmetric—the central finding.

### Contributions

(1) *Transfer is directional*. On architecture-matched models with known training data, structural transfer between language and biology is strongly asymmetric—a language model moved to biology retains far more competence (off-domain drop 0.08) than a biological model moved to language (0.36). (2) *Scaling amplifies the asymmetry*. Language→biology transfer strengthens with model size, whereas biology→language transfer *decays* toward chance; scaling does not universally improve transfer but increases domain specialization—a pattern reproduced across two independent protein-model families. (3) *Structural similarity does not imply transferable intelligence*. Language and biological sequences share structural regularities, yet this shared structure does not license symmetric transfer—a principle for cross-domain foundation-model learning beyond this particular pair of domains.

## 2 A unified protocol for measuring transfer in both directions

All experiments share one schema: a structural pair (*x*_1_, *x*_2_) with a binary label indicating whether the pair is a structural match (paraphrase / homolog) or mismatch (adversarial non-paraphrase / non-homolog). Natural-language pairs are drawn from PAWS-X (English), which contrasts sentences with near-identical lexical content but different word order; biological pairs are protein-sequence pairs labelled homologous vs. non-homologous. A model trained on one domain is evaluated zero-shot on the other, so the two transfer directions are measured with identical code, identical model class, and identical metrics. Because a binary head may adopt either polarity, we report flip-best accuracy against the majority-class baseline and, throughout, audit raw accuracy and seed distributions rather than a single run (Methods). Crucially, the central comparisons use models whose training corpora are fully known (GPT-2, BERT, ESM-2, ProtBERT), so measured transfer cannot be attributed to biological data leaked during pretraining.

## 3 Part I: Forward transfer (language → biology)

Under this protocol we first confirm the forward effect. A GPT-2 classifier fine-tuned only on PAWS-X and evaluated zero-shot on protein-homology pairs transfers well above chance: in the bidirectional protocol (Table 1) forward transfer reaches 0.68 *±* 0.10 flip-best accuracy across 15 seeds (protein majority 0.50), reproducing prior reports of ∼0.80–0.84 and confirming that natural-language structural training instills a capability that carries into biological-sequence discrimination. This forward result is not the contribution of the present work—it is the first of two pieces of evidence whose *comparison* is our subject. We therefore report it under controlled conditions (known training data, multiple seeds, explicit polarity handling) and turn to the reverse direction.

**Table 1:** Bidirectional transfer under one protocol (GPT-2). Forward transfer far exceeds its chance baseline; reverse transfer sits at the majority-class rate. Flip-best accuracy, multiple seeds.

| Direction | Train on | Zero-shot eval | Transfer (flip-best) |  |
| --- | --- | --- | --- | --- |
| Language → Biology | PAWS-X | protein homology | $0.68 \pm 0.10$ | (chance 0.50) |
| Biology → Language | protein homology | PAWS-X | $0.54 \pm 0.01$ | (chance 0.545) |

### The forward effect is structural, not paraphrase-specific

A natural concern is that transfer reflects something peculiar to paraphrase identification rather than natural-language *structure*. To test this we fine-tuned GPT-2 on three distinct structural language sources and evaluated each zero-shot on protein homology (Table 2): PAWS-X (paraphrase / word-order), CoLA (grammatical acceptability), and a synthetic Dyck nested-bracket task (pure hierarchical structure, no semantics). All three transfer well above chance and to a comparable degree (AUC 0.74–0.77). That even a purely syntactic bracket-matching task transfers to protein homology indicates the forward effect is a general consequence of learning natural-language structure, not an artifact of the paraphrase task.

**Table 2:** Forward transfer is structural, not paraphrase-specific. GPT-2 fine-tuned on three distinct natural-language structural tasks, evaluated zero-shot on protein homology (mean *±* s.d., 5 seeds). All transfer comparably above chance.

| Language source (fine-tuning) | Protein AUC | Transfer (flip-best) |
| --- | --- | --- |
| PAWS-X (paraphrase / word order) | $0.77 \pm 0.13$ | $0.68 \pm 0.10$ |
| CoLA (grammatical acceptability) | $0.76 \pm 0.10$ | $0.67 \pm 0.03$ |
| Dyck (nested brackets, pure structure) | $0.74 \pm 0.15$ | $0.64 \pm 0.04$ |

### Why we do not rely on large commercial models

Forward transfer has also been reported to strengthen dramatically at scale: instruction-tuned and frontier models (e.g. Llama-3.1, Qwen-3, Gemini) reach near-perfect accuracy on standard homology and ∼75% on remote homology, with chain-of-thought traces that appear to reason about structural motifs [24, 2]. We deliberately do *not* build our claims on these results. The training corpora of such models are undisclosed and almost certainly contain biological databases (UniProt, Swiss-Prot, SCOP) and biology text, so their apparent structural reasoning cannot be distinguished from memorization or contamination—indeed their chain-of-thought often reveals explicit prior knowledge of protein families. This identifiability problem is precisely why our analysis is confined to models whose training data is fully known (GPT-2, BERT, ESM-2, ProtBERT); it is also what makes the directionality we report interpretable rather than an artifact of undocumented pretraining.

## 4 Part II: Reverse transfer (biology → language) is weak

We evaluate the reverse direction with progressively stronger methods; none yields robust transfer.

### Fine-tuning (100 seeds)

A GPT-2 classifier trained on protein homology learns the biological task nearly perfectly (in-domain accuracy 0.99) but transfers to PAWS-X at 0.543 *±* 0.008, indistinguishable from both the untrained baseline (0.544) and the majority-class rate (0.5465). Across 100 random seeds, *zero* exceed the baseline by more than 0.05—in sharp contrast to the forward direction, which exhibits a high-transfer tail.

### Continued pretraining with matched-token controls

To test whether transfer emerges when language ability is *preserved* rather than overwritten, we continue-pretrain GPT-2 (next-token objective) on 50M tokens of real protein, and on three matched-token controls: shuffled protein (same amino-acid composition, destroyed structure), random amino acids, and English text. We then measure sample-efficiency on PAWS-X (Table 3). At 124M parameters, real protein beats its shuffled control by +0.09 to +0.11 in the low-resource regime—a genuine structural signal—but it does *not* beat text-CPT, and the effect *vanishes* at 355M (protein−shuffled ≈ 0) as the task saturates. On an adversarial nested-bracket (Dyck) task—designed, in the spirit of adversarial diagnostics such as HANS [25], to resist shallow heuristics and remain hard at 355M—protein-CPT shows no advantage over its shuffled control at either scale. Reverse structural benefit is thus real only in a narrow, low-capacity, low-difficulty corner, and is always dominated by natural-language pretraining.

**Table 3:** Iso-token continued-pretraining ablation, PAWS-X sample-efficiency. Real protein beats its shuffled control only at 124M in the low-data regime, never beats text, and the effect disappears at 355M. Mean over 3 seeds.

| Scale | Data frac | base | text | protein | shuffled | $\Delta(\text{prot}-\text{shuf})$ |
| --- | --- | --- | --- | --- | --- | --- |
| 124M | 5% | 0.604 | 0.698 | 0.645 | 0.559 | +0.087 |
| 124M | 10% | 0.777 | 0.802 | 0.789 | 0.675 | +0.114 |
| 355M | 5% | 0.826 | 0.839 | 0.832 | 0.831 | +0.001 |
| 355M | 10% | 0.878 | 0.879 | 0.884 | 0.884 | +0.001 |

### Biological foundation models

We test whether purpose-built protein language models [17, 18] transfer to language. Fine-tuned on PAWS-X, ESM-2 (8M) reaches 0.641—above the 0.545 majority—but this signal is purely structural: it collapses to chance on grammatical acceptability (CoLA, MCC ≈ 0) and entailment (RTE), tasks requiring lexical/semantic access that ESM-2’s 33-token amino-acid vocabulary cannot provide (69% of English becomes `<unk>`). Critically, the signal does not scale: across the ESM-2 ladder (8M → 650M) PAWS-X accuracy *decays* (0.641 → 0.569), and ProtBERT (420M, a different family and architecture) lands at exactly 0.569—the identical majority-class collapse. Two independent large protein models converge on chance, establishing this as a general property of protein LMs rather than a quirk of one model.

### Scope

Symmetry with the forward direction would ideally also test very large models in the reverse direction. This is, however, both harder and less interpretable: frontier biological or general models are typically closed-weight, precluding the iso-token continued-pretraining and matched-control ablations that anchor our reverse analysis, and their pretraining already includes vast natural-language corpora—a reverse “contamination” that would confound any apparent biology→language transfer. We therefore evaluate the reverse direction, as the forward, on open models with documented training data (ESM-2, ProtBERT), which also carry no natural-language pretraining and thus provide a clean test. The consistency of the effect across two families and four scales indicates the conclusion is not an artifact of small scale.

## 5 Part III: The transfer is directional

Placing both directions on architecture-matched models with known training data quantifies the asymmetry directly (Table 4). A pure protein model (ESM-2) solves biology (0.999) but barely exceeds chance on language (0.641 vs. 0.545): an off-domain drop of 0.36. A size-matched English model (BERT) solves language (0.732) and, moved to biology, retains real competence (0.656 vs. 0.50 chance): an off-domain drop of only 0.08. The language model transfers roughly four times more of its competence across the domain boundary than the biological model does. Because these are small models with fully documented training corpora, this asymmetry cannot be an artifact of biological pretraining contamination—the principal confound in large-model studies.

**Table 4:** **Architecture-matched 2 *×* 2 cross-modal transfer** (accuracy, 3 seeds; known-training-data models, no contamination confound). A language model retains far more off-domain competence than a biological model.

| Backbone | Pretraining | → protein homology | → PAWS-X |
| --- | --- | --- | --- |
| ESM-2 (8M) | protein | **0.999** | 0.641 |
| BERT-small | English | 0.656 | **0.732** |
| BERT-tiny | English | 0.636 | 0.583 |
| off-domain drop from home task |  | bio-model: 0.36 | language-model: 0.08 |

Scaling widens the gap rather than closing it. English pretraining transfers *more* with size (BERT-tiny→base: 0.583 → 0.911 on PAWS-X), while protein pretraining transfers *less* with size (ESM 0.641 → 0.569), converging to chance (Fig. 2). Directionality is therefore not a small-model accident but a stable, scale-amplified, cross-family property.

**Figure 2:**
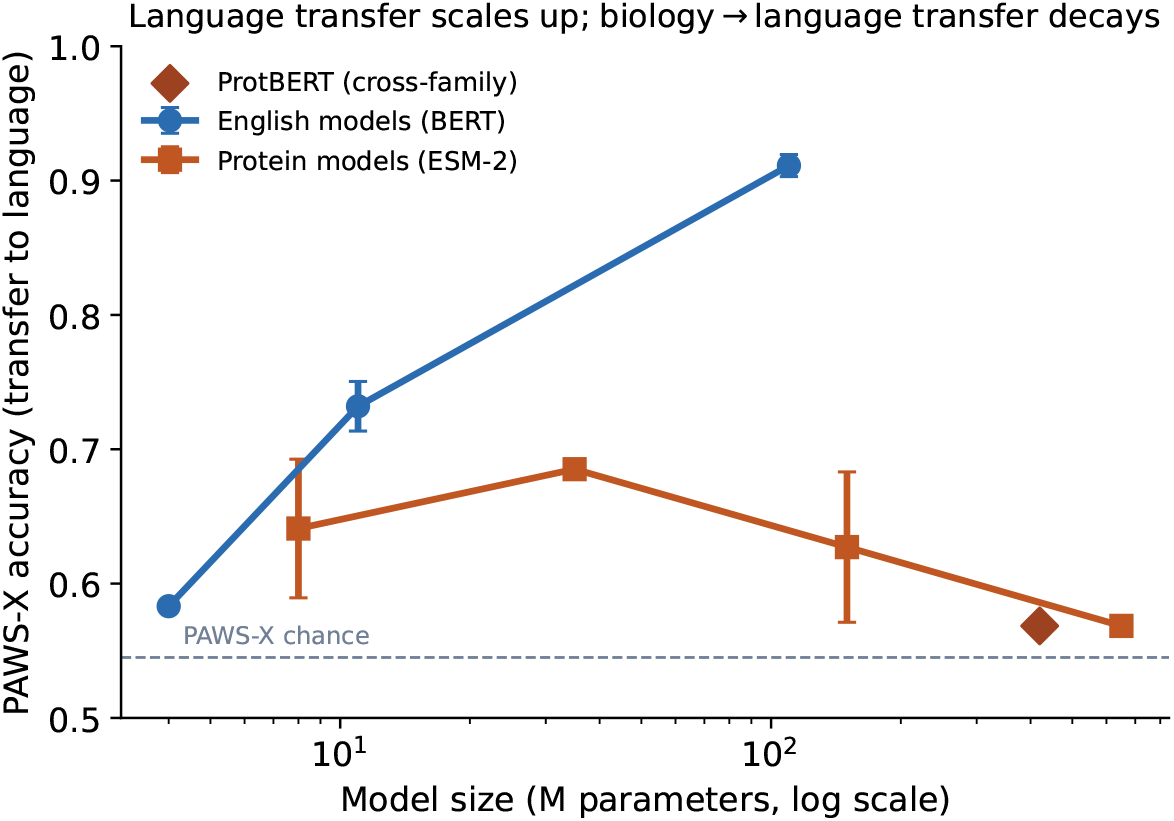
Scaling widens the asymmetry. Transfer to natural language (PAWS-X accuracy) versus model size. English pretraining transfers *more* with scale (BERT, blue), whereas protein pretraining transfers *less*, decaying toward chance (ESM-2, orange); ProtBERT (a different family, red diamond) confirms the collapse at scale. Mean *±* s.d., 3 seeds.

## 6 Part IV: Mechanistic origin of the asymmetry

Three analyses localize the source of the asymmetry. They rule out two intuitive explanations (static representational sharing; an induced difference-detecting circuit) and point instead to a difference in *learnability* : the target capability emerges during source-domain training in the forward direction only.

### Representations are not globally aligned (M1)

We first asked whether the two domains occupy a shared representational subspace by computing linear CKA [26, 27] between a model’s per-layer representations of PAWS-X pairs and of protein pairs, following the representation-analysis and probing tradition [28, 29]. Across base GPT-2, text-continued and protein-continued models, CKA remains at the row-shuffled control floor (≈ 0.04–0.05) at every layer (Table 5). The domains do *not* share a globally aligned geometry; the transfer that does occur cannot be explained by a static shared manifold.

**Table 5:**
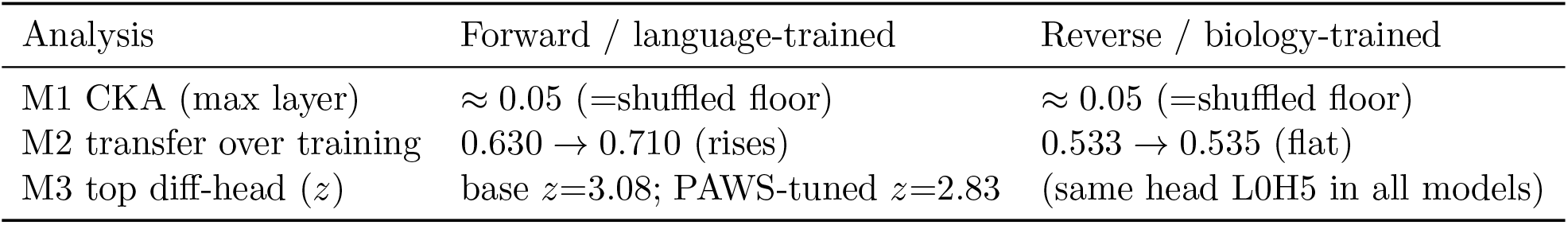
Mechanistic analyses. (M1) cross-modal CKA at the shuffled floor; (M2) transfer emerges only in the forward direction; (M3) protein training does not induce NL difference heads.

### Transfer emerges during forward training but never during reverse training (M2)

We snapshotted zero-shot transfer accuracy every 60 optimization steps. In the forward direction (Fig. 3), transfer to protein *rises monotonically* as the model learns the language task (0.630 →0.710, peak 0.773; +0.08 over training). In the reverse direction, transfer to language stays pinned at chance throughout (0.533 → 0.535; +0.001). Forward transfer is an emergent by-product of learning language structure; reverse transfer never emerges, at any point in training.

**Figure 3:**
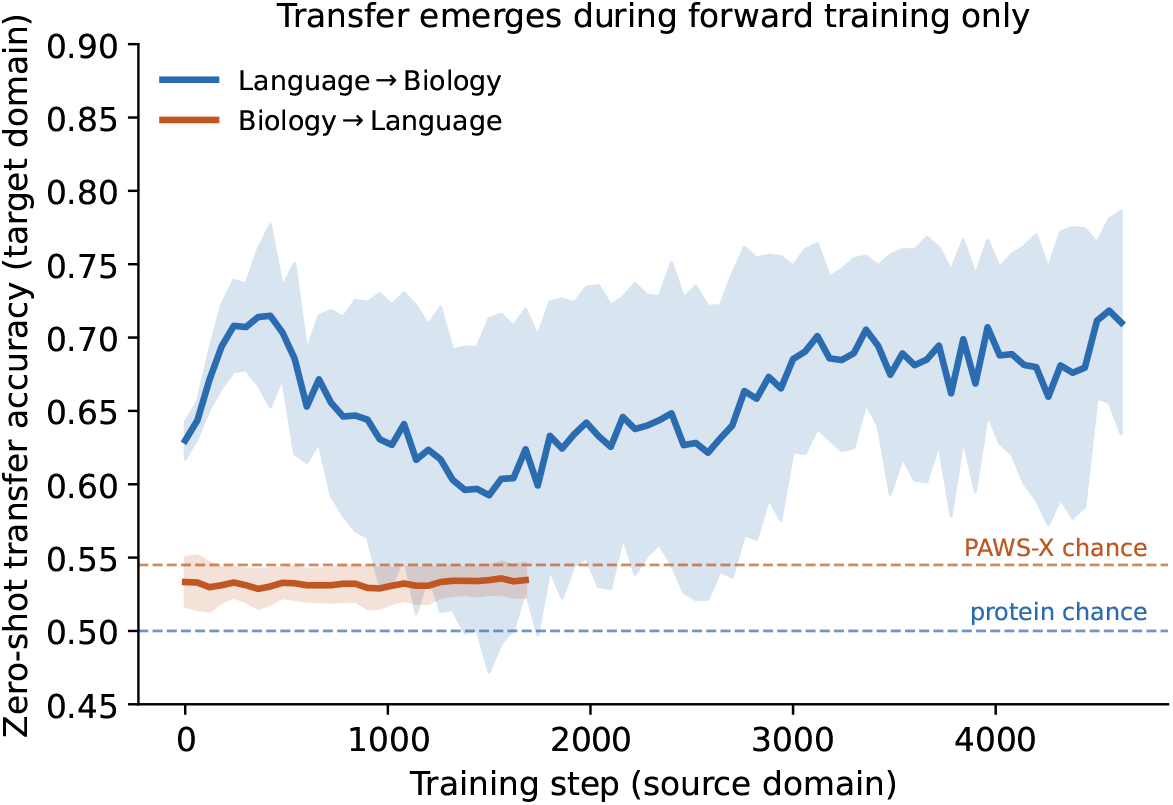
Transfer emerges during forward training only (M2). Zero-shot transfer accuracy on the target domain, snapshotted every 60 steps during source-domain training (mean *±* s.d., 3 seeds). Language →biology transfer rises steadily as the model learns language structure; biology →language transfer remains at the chance line throughout training.

### Difference-detecting heads are not the mechanism (M3)

A natural candidate mechanism, proposed in prior work, is that structural training induces attention heads acting as “difference operators” that localize the positions differing between a pair. We tested this rigorously with three variants: whether protein training induces heads that mark NL differences, whether language training induces heads that mark protein mutations, and—most carefully—a head-level difference-selectivity score (attention flowing into differing versus matched positions) *z*-scored against each model’s own head distribution. The claim does not survive. Difference-selective heads *exist* but are *pretraining artifacts* : base GPT-2 already contains the strongest NL difference head (layer 0 head 5, *z* = 3.08) before any fine-tuning, and neither PAWS training (*z* = 2.83, if anything lower) nor protein training strengthens it; the identical head tops all model variants (Table 5). The asymmetry is therefore *not* explained by differential induction of difference heads. We report this as a cautious correction: a mechanism attractive on qualitative single-head inspection does not hold under systematic, baseline-controlled testing.

### Interpretation

The three analyses locate the asymmetry precisely. It is not a matter of static representational sharing (M1: the domains are not even globally aligned) nor of a specific induced circuit (M3: difference heads are pretraining artifacts, not trained-in operators). What distinguishes the two directions is *whether the target capability emerges as a by-product of source-domain learning* (M2): learning language structure progressively confers biological discrimination, whereas learning biological structure never confers linguistic discrimination at any point in training. We therefore frame the directionality as a difference in the *learnability* of cross-domain transfer, and *hypothesize*— as an account consistent with, but not proven by, the present evidence—that natural-language training, with its demands of abstraction, recursion and compositionality, shapes representations that generalize more readily to other structured domains than do the evolution-, folding- and fitness-shaped representations of protein models. We explicitly do not claim a specific mechanistic circuit; our difference-head analysis shows that an intuitively appealing candidate does not, in fact, account for the effect.

## 7 Discussion

Our results establish that the existence of shared structural regularities between natural language and biological sequences does *not* imply symmetric representational transfer, highlighting an intrinsic directionality in cross-domain foundation-model learning. This reframes the much-discussed language→biology transfer: it is not evidence of a symmetric universal grammar but of an *asymmetric* one, in which language serves as the stronger prior. The directionality is stable across fine-tuning, continued pretraining, scaling, model families, and task difficulty, and—because it is demonstrated on models with known training data—is not attributable to pretraining contamination. Our mechanistic analyses rule out two intuitive explanations—static representational alignment and an induced difference-detecting circuit—and localize the asymmetry to learnability: biological discrimination emerges as a by-product of learning language structure, but not the converse. We frame the deeper account as a hypothesis to be tested rather than a settled claim: the abstraction, recursion and compositionality demanded by natural language may shape representations that generalize to other structured domains more readily than the evolution-, folding- and fitness-shaped representations of biological models.

A striking corollary concerns scale. It is tempting to assume that whatever a small model fails to transfer, a larger one will; our scaling analysis shows the opposite for the reverse direction, where transfer *decays* toward chance as protein models grow. Scaling does not universally improve cross-domain transfer—it can instead deepen domain specialization—and the two directions diverge rather than converge. More broadly, our findings caution against inferring transferable capability from structural resemblance: two domains can share statistical and structural regularities, and a model can be highly competent in each, without transfer between them being symmetric or even present. We suggest that the direction and scaling behaviour of cross-domain transfer, rather than the mere existence of shared structure, is the more informative probe of what foundation models actually learn.

## Methods

### Task schema and datasets

Every task is a binary structural-pair classification: given (*x*_1_, *x*_2_), predict match (label 1) vs. mismatch (label 0). Natural-language pairs use PAWS-X English [30] (49,401 train, 2,000 test), whose adversarial pairs share lexical content but differ in word order. Biological pairs use a balanced protein-homology set: positives are Swiss-Prot pairs with BLASTp *E*-value ≤ 10^−10^ and sequence identity above threshold; negatives are length-matched random pairs verified non-homologous by global alignment; 20,000 balanced pairs, lengths 40–250 residues. Grammatical-acceptability (CoLA) and entailment (RTE) tasks use the GLUE distributions [31]. The exact dataset files, splits, and accession-level metadata are released (Data availability).

### Models

Language backbones: GPT-2 (124M) and GPT-2-medium (355M) [32]; BERT-tiny (4M), BERT-small (11M), BERT-base (110M) [33]. Biological backbones: ESM-2 at 8M/35M/150M/650M [15] and ProtBERT (420M) [19]. All are public checkpoints with documented training corpora, so no biological data could have leaked into the language models and no natural-language data into the protein models—this is what makes the 2*×*2 comparison free of the contamination confound.

### Fine-tuning and zero-shot transfer

For classification transfer we attach a linear head and fine-tune on the source domain (AdamW, learning rate 1*×*10^−5^, batch size 32, 3–4 epochs, constant-with-warmup schedule), then evaluate zero-shot on the target domain. Because a freshly initialized binary head has arbitrary polarity, we report flip-best accuracy max(acc, 1 − acc) against the majority-class baseline, and additionally report raw accuracy and full per-seed distributions; we never select polarity to inflate a headline number. Forward and reverse directions are run by the same script with source and target swapped.

### Iso-token continued pretraining

From GPT-2 we continue-pretrain (causal-LM objective, block size 512) on exactly 5.0 *×* 10^7^ tokens per corpus, matched across four groups so that only sequence *content* differs: (G2) real Swiss-Prot proteins rendered as space-separated residues `<protein> M K V … </protein>`; (G3) the same sequences with residues shuffled within each protein (identical amino-acid frequency and length, destroyed structure); (G4) residues resampled i.i.d. from the background amino-acid distribution; (G1) OpenWebText English. Token counts are measured with the GPT-2 tokenizer to guarantee the iso-token constraint. All groups use identical optimizer, steps (≈3050), and batch size. Sample-efficiency is then measured by fine-tuning each resulting checkpoint on *{*1, 5, 10, 100*}*% of a downstream task.

### Hard structural control (Dyck)

To obtain a natural-language-side structural task that does not saturate at 355M, we generate Dyck-language nested-bracket sequences over three bracket types; positives are well-formed, negatives are corrupted by a single bracket edit at a random depth, so local heuristics fail and only tracking the nesting stack succeeds. Difficulty is tuned by length (*L* ∈ *{*40, 60*}*).

### Mechanistic analyses

(M1) Representational alignment: linear CKA [26] between a model’s per-layer last-token representations of PAWS-X pairs and of protein pairs, against a row-shuffled floor. (M2) Learning dynamics: during training we snapshot zero-shot transfer accuracy on a fixed held-out target set every 60 steps, yielding a transfer-vs-step trajectory for each direction. (M3) Difference-operator heads: for each attention head we measure a difference-selectivity score—attention flowing into the positions that differ between *x*_1_ and *x*_2_ minus that into matched positions—*z*-scored against the same model’s head distribution, and compare base GPT-2, PAWS-tuned, and protein-continued models on both modalities to test whether either training direction *induces* difference-detecting heads above the pretrained baseline.

### Statistics

Distribution experiments use up to 100 random seeds; all other experiments use 3 seeds and report mean *±* s.d. Chance levels are the majority-class rates (protein 0.50, PAWS-X 0.545).

## Data availability

Datasets are released at https://huggingface.co/dnagpt/biopaws, including exact file names, splits, and accession metadata.

## Code availability

All code for corpus construction, continued pretraining, evaluation, and mechanistic analysis is available at https://github.com/maris205/bio2nl.

